# An evaluation of clustering and assembly strategies from Iso-Seq data in the absence of reference genomes in non-model animals

**DOI:** 10.1101/2025.09.18.677004

**Authors:** Klara Eleftheriadi, Marçal Vázquez-Valls, Rosa Fernández

**Affiliations:** Metazoa Phylogenomics and Genome Evolution Lab, Institute of Evolutionary Biology (CSIC-UPF), Barcelona (Spain)

**Keywords:** Long-read sequencing, PacBio IsoSeq, de-novo transcriptome assemblies, non-model organisms

## Abstract

Transcriptome assembly enables the recovery of expressed genes and isoforms, but the optimal strategy for reconstructing transcriptomes from long-read sequencing remains unresolved. In particular, establishing best practices for generating accurate gene models and selecting representative isoforms is essential for comparative genomics, since orthology inference typically requires only the longest isoform per gene model. Here, we systematically compare clustering and de novo assembly methods using PacBio Iso-Seq data from diverse invertebrate lineages with the goal of identifying the most optimal methodology for isoform selection in the absence of dedicated pipelines. We evaluate four approaches: IsoSeq3 (isoseq3 cluster), CD-HIT, RNA-Bloom2 and isONform, all benchmarked against short-read Trinity assemblies. Assembly quality was assessed using BUSCO completeness, short-read mapping rates, coding sequence recovery, longest isoform prediction, and SQANTI3 structural classification. Our results show that CD-HIT clustering at high similarity thresholds (≥99%) yields the most complete and coding-rich long-read transcriptomes, rivaling Trinity while avoiding its high redundancy. SQANTI3 classification further confirms that CD-HIT 99 recovers the highest number of full-splice-match transcripts among all methods. Consensus-based methods such as IsoSeq3 and isONform recover fewer single-copy orthologs (mirrored in a lower BUSCO score) and achieve lower mapping rates, while RNA-Bloom2 provides intermediate performance with reduced duplication. Together, these findings establish, to date, CD-HIT as a robust and practical strategy for transcriptome reconstruction from long-read data when genomic references are unavailable. This work provides practical guidance for deriving high-quality gene models and selecting representative isoforms for orthology inference in non-model species.

## Introduction

Transcriptome assembly is the computational reconstruction of all RNA transcripts present in a cell, tissue, or organism. These assemblies provide insight into gene expression, alternative splicing and regulatory networks that cannot be captured by genomic information alone. Thus, transcriptome assembly has become an essential tool in fields ranging from developmental biology and evolutionary genomics to agriculture and medicine, where it enables the discovery of novel genes, non-coding RNAs, and biomarkers relevant to health and disease (Wang et al. 2009; Conesa et al. 2016).

Broadly, there are two main categories for reconstructing a transcriptome assembly from raw reads: genome-guided and de novo. Genome-guided methods leverage an available reference genome to align the reads and assemble the transcripts, which typically improves accuracy, but their utility is limited in species with incomplete or poorly annotated references (Haas et al. 2013). In contrast, de novo assemblers reconstruct transcripts directly from the sequencing data, making them indispensable for non-model organisms and for identifying transcriptomic features absent from reference genomes. However, de novo approaches are computationally more demanding and may be prone to artifacts such as fragmented or chimeric transcripts (Smith-Unna et al. 2016). Both approaches involve trade-offs that must be carefully considered depending on the organism and resources available.

Short-read sequencing platforms such as Illumina have dominated the field for more than a decade because of their cost-effectiveness and high throughput, and they remain widely used in transcriptomic research. Nevertheless, the short length of reads prevents the direct recovery of full-length transcripts and isoforms, limiting the completeness and accuracy of transcriptomes (Byrne et al. 2019). Reliance on fragmentary data introduces particular challenges in reconstructing complex transcriptomes with extensive alternative splicing, often resulting in incomplete, fragmented, or chimeric assemblies (Smith-Unna et al. 2016).

The advent of third-generation sequencing technologies, particularly Pacific Biosciences’ (PacBio) single-molecule real-time (SMRT) Iso-Seq protocol, has marked a turning point in transcriptomics. PacBio high-fidelity (HiFi) sequencing now produces long, accurate reads (>99.9% median accuracy), frequently spanning tens of kilobases and preserving exon order and orientation (Gonzalez-Garay 2016; Wijeratne et al. 2024; Wang et al. 2025). These properties improve isoform detection and reduce the complexity of transcriptome reconstruction.

In eukaryotes, a single gene typically contains multiple exons separated by introns, and through processes such as alternative splicing, it can give rise to multiple transcript isoforms that together define the gene model. When a high-quality reference genome is available, gene models are inferred with confidence by mapping long-read transcripts to genomic coordinates using established pipelines (e.g. StringTie, Mandalorion) (Kovaka et al. 2019; Volden et al. 2023). In the absence of a genome, gene-level inference must be approximated directly from the transcript sequences. In short-read RNA-seq, where the read-length does not span full transcripts, graph-based assembly approaches (e.g. Trinity) (Grabherr et al. 2011) are typically used to first reconstruct isoforms before aggregating them into putative gene-level clusters, although these methods suffer from fragmentation and redundancy. By contrast, long-read sequencing technologies capture full-length transcripts directly, reducing the need for computational reconstruction of transcript sequences (Sahlin and Medvedev 2020). Nonetheless, grouping these transcript isoforms into clusters to construct gene models remains a major challenge (Sahlin and Medvedev 2020). Many downstream analyses in comparative studies, including orthology inference (Li et al. 2003; Emms and Kelly 2019), phylogenomic analyses (Fernández et al. 2014; Fernández and Gabaldón 2020; Balart-García et al. 2021), gene-level expression quantification (Davidson and Oshlack 2014), require transcript isoforms to be collapsed into putative gene models. This is typically achieved by retaining a single representative isoform per gene, most often the longest coding sequence.

However, the optimal strategy for reconstructing de novo transcriptome assemblies with gene models from Iso-Seq data remains unclear. A key question persists: Is clustering alone sufficient to recover a complete transcriptome, or does a de novo assembly step still offer advantages, even with long-read data? While many studies default to the PacBio Iso-Seq clustering pipeline, few have rigorously assessed its completeness or quality, especially in non-model species where benchmarking remains sparse (Pootakham et al. 2020; Ali et al. 2021). Existing benchmarking efforts have largely focused on model organisms or differential expression, and have not been systematically evaluated across multiple lineages (Yan et al. 2026). To our knowledge, no study has simultaneously evaluated these approaches across taxonomically diverse non-model lineages.

In this study, we address this gap by systematically comparing clustering algorithms and de novo assembly methods for long-read transcriptome reconstruction using experimental data from 17 invertebrate lineages. We evaluate the performance of four tools, categorized as (i) clustering-based tools, including PacBio’s isoseq3 cluster module (IsoSeq3) (https://github.com/PacificBiosciences/pbbioconda) and CD-HIT (Fu et al. 2012), and (ii) graph-based de novo assemblers, including RNA-Bloom2 (Nip et al. 2020) and isONform (Petri and Sahlin 2023). Additionally, we use Trinity (Grabherr et al. 2011) as a benchmark for completeness and comparison with long-read approaches. To assess transcriptome quality, we employ multiple metrics: BUSCO scores (Simão et al. 2015), short-read alignment rates, open reading frame (ORF) recovery, the total number of longest protein isoforms per assembly and SQANTI3 structural classification of transcript splice junctions (Pardo-Palacios et al. 2024) leveraging the highly-curated reference genome of *Caenorhabditis elegans*. Through this comprehensive comparison, we aim to identify the most robust and practical strategy, to date, for constructing high-quality transcriptomes from long-read Iso-Seq data, particularly in non-model organisms, and provide practical recommendations for the biodiversity genomics community.

## Materials and Methods

### Sampling and sequencing

The dataset analyzed in this study was generated previously by (Martínez-Redondo et al. 2025) and comprises 17 species across 7 animal phyla (Figure 1), including both model organisms such as *Caenorhabditis elegans* and *Schmidtea mediterranea*, as well as numerous non-model species from diverse ecological and evolutionary lineages. This broad taxonomic sampling provides a robust framework for evaluating transcriptome assembly methods across deep evolutionary divergence.

**Figure 1.**
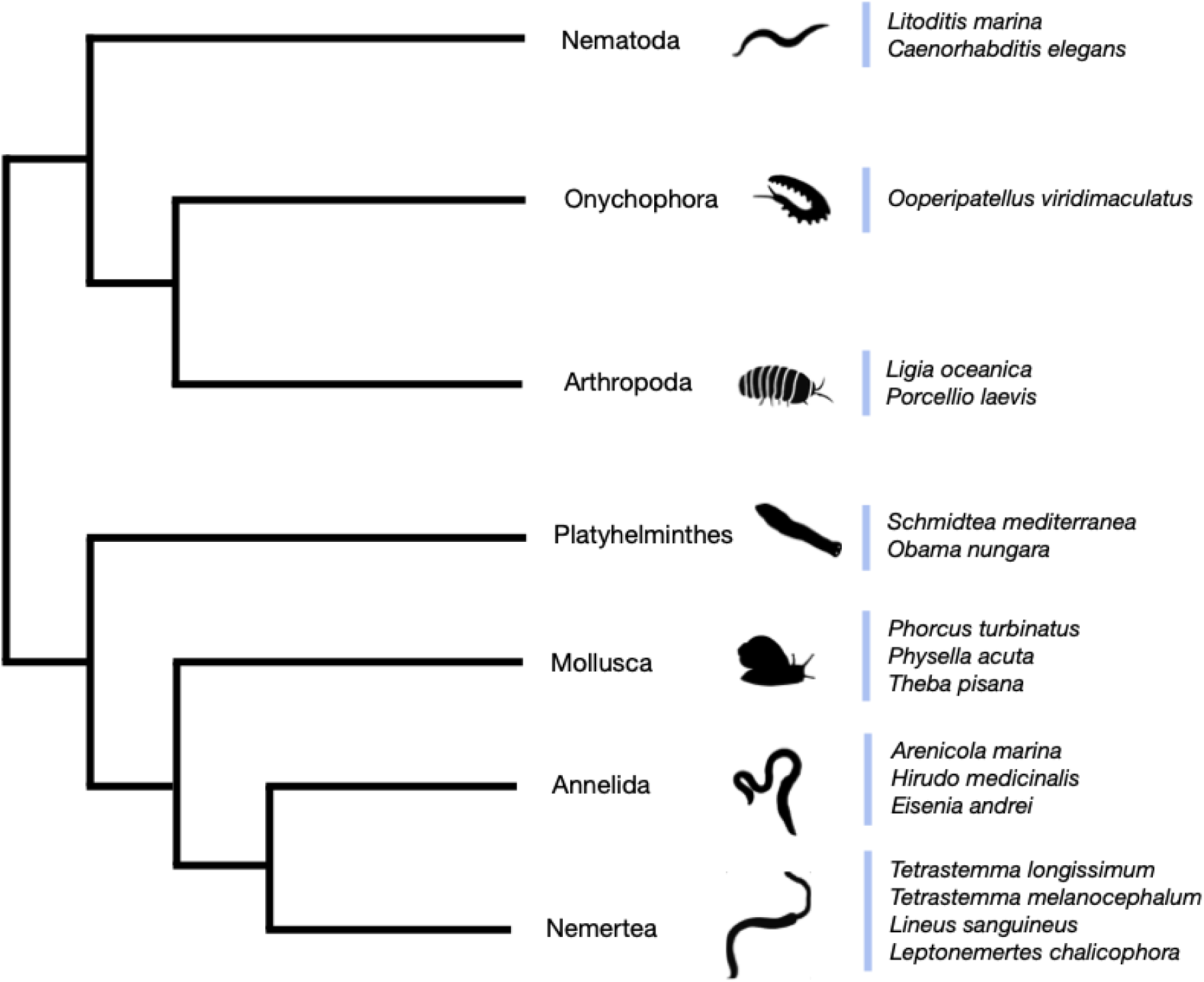
Phylogenetic distribution of sampled species. Phylogenetic tree showing the 17 animal species used in this study, spanning seven metazoan phyla: Arthropoda, Annelida, Onychophora, Mollusca, Nemertea, Platyhelminthes, and Nematoda. Species names are listed alongside their respective phyla.

For each species, both long-read Iso-Seq data and short-read RNA-Seq data were generated as described in (Martínez-Redondo et al. 2025). Briefly, to maximize transcript diversity, between 35 and 60 individuals were exposed to a range of experimental treatments designed to induce diverse transcriptional responses prior to RNA-extraction. RNA extractions after stress experiments were performed using the TRIzol® reagent (Invitrogen, USA) method following the manufacturer’s instructions and using MaXtract® High Density tubes (Qiagen) to minimize DNA contamination prior to mechanical sample homogenization either by plastic rotor pestles or by ceramic mortar, depending on the sample. The concentration of all samples was assessed by Qubit RNA BR Assay kit (Thermo Fisher Scientific). Libraries were prepared with the TruSeq Stranded mRNA library preparation kit (Illumina), and sequenced on a NovaSeq 6000 (Illumina, 2 × 150 bp) for a minimum of 6Gb coverage.

The same RNA extractions used for short-read sequencing were pooled together in each species to prepare an Iso-Seq library. RNA samples were subjected to DNAse treatment using the Turbo DNA-free DNase (Invitrogen) following the manufacturer’s instructions. SMRTbell libraries were generated following the procedure ‘Preparing Iso-Seq® libraries using SMRTbell® prep kit 3.0 (PN 102-396-000 REV02 APR2022)’. To enable pooling of multiple samples on 1 SMRTcell, libraries were made using barcoded adapters. The libraries were sequenced on a Sequel-IIe using Sequel II sequencing kit 2.0 and Binding kit 3.1 with 24 hr movie-time.

The majority of the species included in this dataset are non-model organisms that are undersampled and poorly characterized both at the genomic and transcriptomic level. This taxonomically diverse and biologically relevant dataset provides a robust framework for evaluating transcriptome reconstruction strategies across different assembly approaches and sequencing technologies.

### Long and short reads preprocessing

Prior to the construction of de novo transcriptome assemblies, we obtained the Full-Length Non-Concatemer (FLNC) reads separately from Iso-Seq data of the 17 species using the preprocessing pipeline available from PacBio. The data were demultiplexed with the Iso-Seq pipeline v4.0.0 (https://github.com/PacificBiosciences/pbbioconda), and cDNA primers were removed using LIMA v2.7.1. PolyA tails and artificial concatemers were removed using isoseq refine v4.0.0.

Raw Illumina RNA-Seq reads were quality controlled with FastQC v0.11.9 (https://www.bioinformatics.babraham.ac.uk/projects/fastqc/) and adapters and low-quality base pairs were removed using Trimmomatic v0.39 (MINLEN: 75, SLIDINGWINDOW: 4:15, LEADING: 10, TRAILING: 10, AVGQUAL: 30) (Bolger et al. 2014). Trimmed RNA-seq reads were quality controlled with FastQC before further analysis.

### Construction of de novo reference transcriptome assemblies using clustering algorithms

We employed two main methods to construct de novo reference transcriptomes using clustering algorithms. The first method was the isoseq3 cluster module (hereafter IsoSeq3) within the PacBio SMRT suite (https://github.com/PacificBiosciences/pbbioconda). Using isoseq3 cluster v4.0.0, we followed the publicly available PacBio protocol for transcriptome construction. In this process, two or more FLNC reads are clustered if they differ by less than 100 bp at the 5′ end and less than 30 bp at the 3′ end, with the additional requirement that there are no internal gaps. The resulting consensus sequences define the transcriptome as High Quality Isoforms, while Low Quality Isoforms are discarded from further analysis.

For the second method, we used CD-HIT v4.8.1 (Li and Godzik 2006), a fast and flexible clustering tool originally developed for clustering large protein sequence databases at various sequence identity thresholds. CD-HIT significantly reduced execution times compared to earlier programs, though it can introduce some redundancy. We used CD-HIT with sequence similarity thresholds of 95%, 96%, 97%, 98%, and 99% to cluster the preprocessed FLNC reads. For each threshold, the longest sequence in each cluster was retained as the cluster representative, forming five distinct reference transcriptomes composed of the most “informative” sequences. While CD-HIT was first designed for protein clustering, its underlying greedy incremental algorithm (Holm and Sander 1998) is also well-suited for DNA and RNA sequence clustering, as implemented in the CD-HIT-EST tool used in our analysis (Li and Godzik 2006).

### Construction of de novo reference transcriptomes using graph-based assembly methods

De novo transcriptome assembly can be achieved through several algorithmic approaches, with graph-based algorithms among the most widely used for reconstructing transcripts from sequencing reads. For this purpose, we chose two widely used software for long-read Iso-Seq data, RNA-Bloom2 (Nip et al. 2020) and isONform (Petri and Sahlin 2023). Then we also used Trinity (Grabherr et al. 2011) to reconstruct assemblies from short Illumina reads for evaluation purposes, as described below.

RNA-Bloom2 is a reference-free assembler tailored for long-read transcriptome sequencing data. We used RNA-Bloom2 v2.0.1 to assemble transcriptomes directly from FLNC reads, without requiring a reference genome. The process takes the FLNC reads as input and produces a final transcriptome assembly as output.

isONform follows a multi-step workflow and operates in conjunction with isONclust. For our assemblies with isONform v0.3.3, we first used isONclust to cluster FLNC reads efficiently, employing a greedy, minimizer-based approach that switches to full alignment as needed (Sahlin and Medvedev 2020). isONform then constructs isoforms from these clusters, generating a set of predicted transcripts in FASTQ format for downstream analysis.

### Quality evaluation of the reference de novo transcriptome assemblies

Trinity is one of the most established reference-free assembly programs for short-read RNA-Seq data (Tzec-Interián et al. 2025). Based on de Bruijn graph algorithms, Trinity enables the de novo assembly of full-length transcripts when a suitable reference genome is unavailable. In this study, we used Trinity v2.11.0 to assemble transcriptomes from short-read Illumina RNA-seq data. These assemblies serve as a key evaluation metric for comparing the performance of the long-read assembly methods.

To compare the quality of transcriptome assemblies produced by different algorithms, we used several global evaluation metrics. Assembly completeness was assessed using BUSCO v5.4.7 (Simão et al. 2015), analyzing the presence of 954 universal single-copy orthologs from the metazoa odb10 database. To evaluate read support, we mapped preprocessed short RNA-seq reads to each reference transcriptome using Minimap2 (Li 2018) and calculated the proportion of mapped reads. Coding sequence recovery was measured with TransDecoder v5.5.0 (https://github.com/TransDecoder/TransDecoder), which identifies candidate coding regions and predicts likely protein sequences in each assembly. Finally, we quantified the number of longest isoforms present in each reconstructed transcriptome, in order to compare the resulting longest protein isoforms per assembly.

To statistically evaluate the performance of each assembler across species for BUSCO completeness and short-read mapping rate, we applied a Friedman test to assess whether the assemblers differed overall, and where significant, we performed pairwise post-hoc comparisons using Wilcoxon signed-rank tests with Holm correction for multiple testing. Assemblers were ranked and ordered in figures by descending mean rank. Statistically homogeneous groups are indicated by compact letter display (CLD; α=0.05).

### Evaluation using a reference genome of the model species *C. elegans*

To further evaluate the structural accuracy of the reconstructed transcriptomes, we performed a reference-guided assessment using a model species included in our dataset, the nematode *Caenorhabditis elegans*, for which a highly curated reference genome assembly and structural annotation are available (GCF_000002985.6). For this purpose, we used SQANTI3 (Pardo-Palacios et al. 2024) which compares transcript models against a reference genome and annotation and classifies the transcripts based on their agreement with the annotated splice junction chains. We focused on four principal categories: full-splice-match (FSM), in which all splice junctions match an annotated transcript; incomplete-splice-match (ISM), in which transcripts match a subset of the splice junctions of an annotated transcript; novel-in-catalog (NIC), containing novel combinations of previously annotated splice junctions; and novel-not-in-catalog (NNC), containing at least one unannotated splice junction.

We applied SQANTI3 QC module to transcriptomes generated by each clustering and assembly strategies, including the raw FLNC, IsoSeq3, isONform, RNA-Bloom2 and CD-HIT at 95-99% sequence similarity thresholds. Transcript sequences were aligned to the *C. elegans* reference genome using minimap2 through the SQANTI3 workflow. In addition, the Illumina short-read RNA-seq data were incorporated as orthogonal evidence by mapping them to the reference genome using STAR v2.7.11b (Dobin et al. 2013). We then compared the counts of the four structural categories among assemblies to evaluate the recovery of the annotated transcript structures.

## Results & Discussion

Here, we compared clustering and assembly strategies for de novo transcriptome reconstruction from long-read Iso-Seq data in 17 predominantly non-model animal species. We evaluated four approaches: (i) IsoSeq3, (ii) CD-HIT clustering at five similarity thresholds (95–99%), (iii) RNA-Bloom2, and (iv) isONform, all of them benchmarked against Trinity assemblies generated from short-read Illumina RNA-seq data, together with the unassembled FLNC reads. In total, 170 transcriptomes were constructed and assessed for quality using BUSCO completeness (metazoa odb10), short-read mapping rates using Minimap2, and the number of predicted protein-coding sequences and longest isoforms per assembly (Figure 2, Table S1).

**Figure 2.**
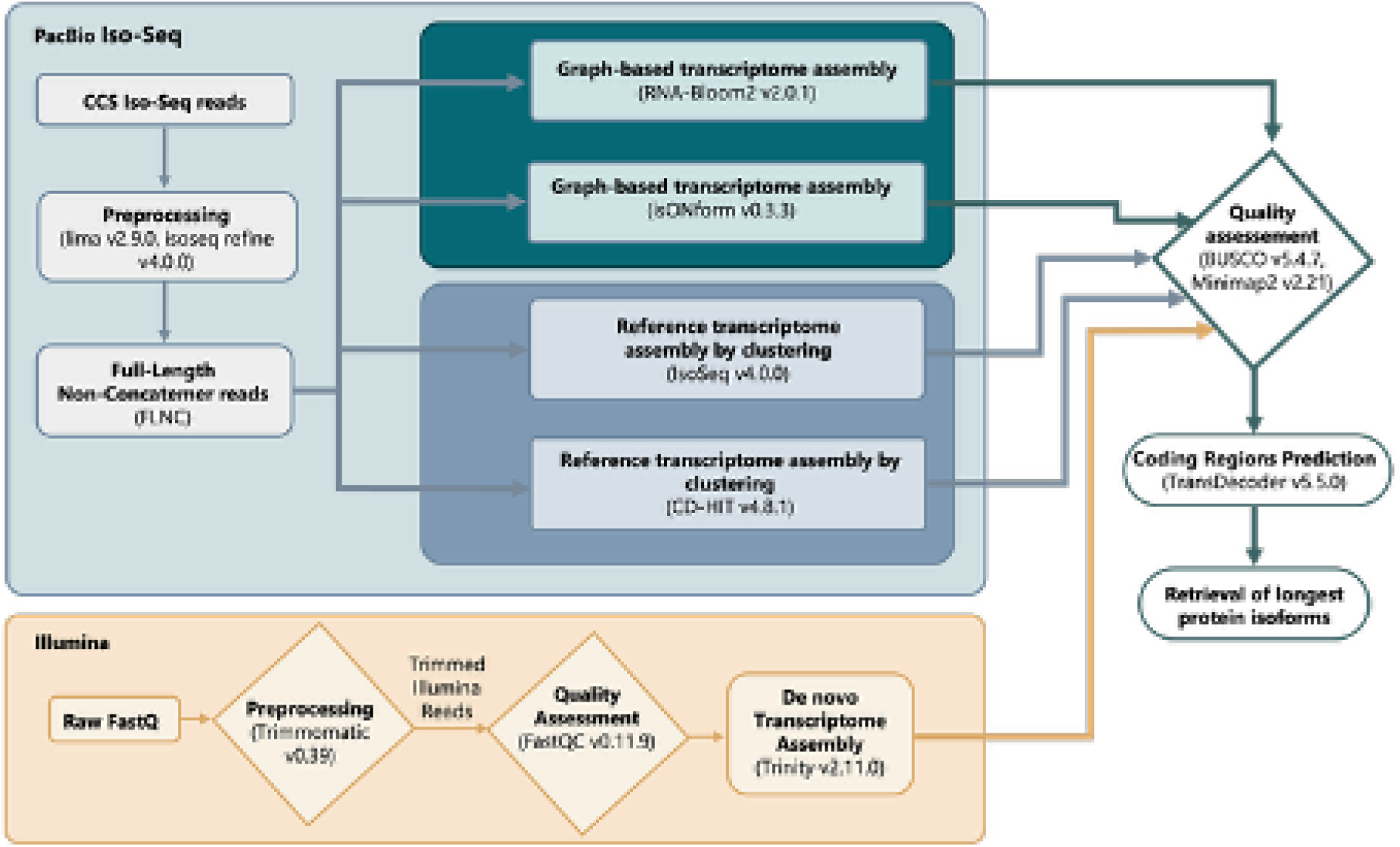
Workflow for evaluating transcriptome assembly strategies. Workflow showing preprocessing, clustering, and assembly of Illumina and PacBio Iso-Seq reads, followed by quality assessment using BUSCO, Minimap2, and TransDecoder to evaluate completeness, read support, and coding potential.

### Assembly completeness

Our results revealed significant differences in BUSCO completeness among assemblers across species. Specifically, CD-HIT at 99% similarity consistently outperformed other long-read methods, with BUSCO scores rivaled only by Trinity (Figure 3a, 4a; Supplementary Figure S1). The higher completeness obtained with CD-HIT likely reflects its use of the longest representative sequence in each cluster, whereas consensus-based approaches such as IsoSeq3 and isONform tend to collapse transcript diversity, potentially discarding biologically valid isoforms or introducing artificial consensus sequences not corresponding to conserved genes.

**Figure 3.**
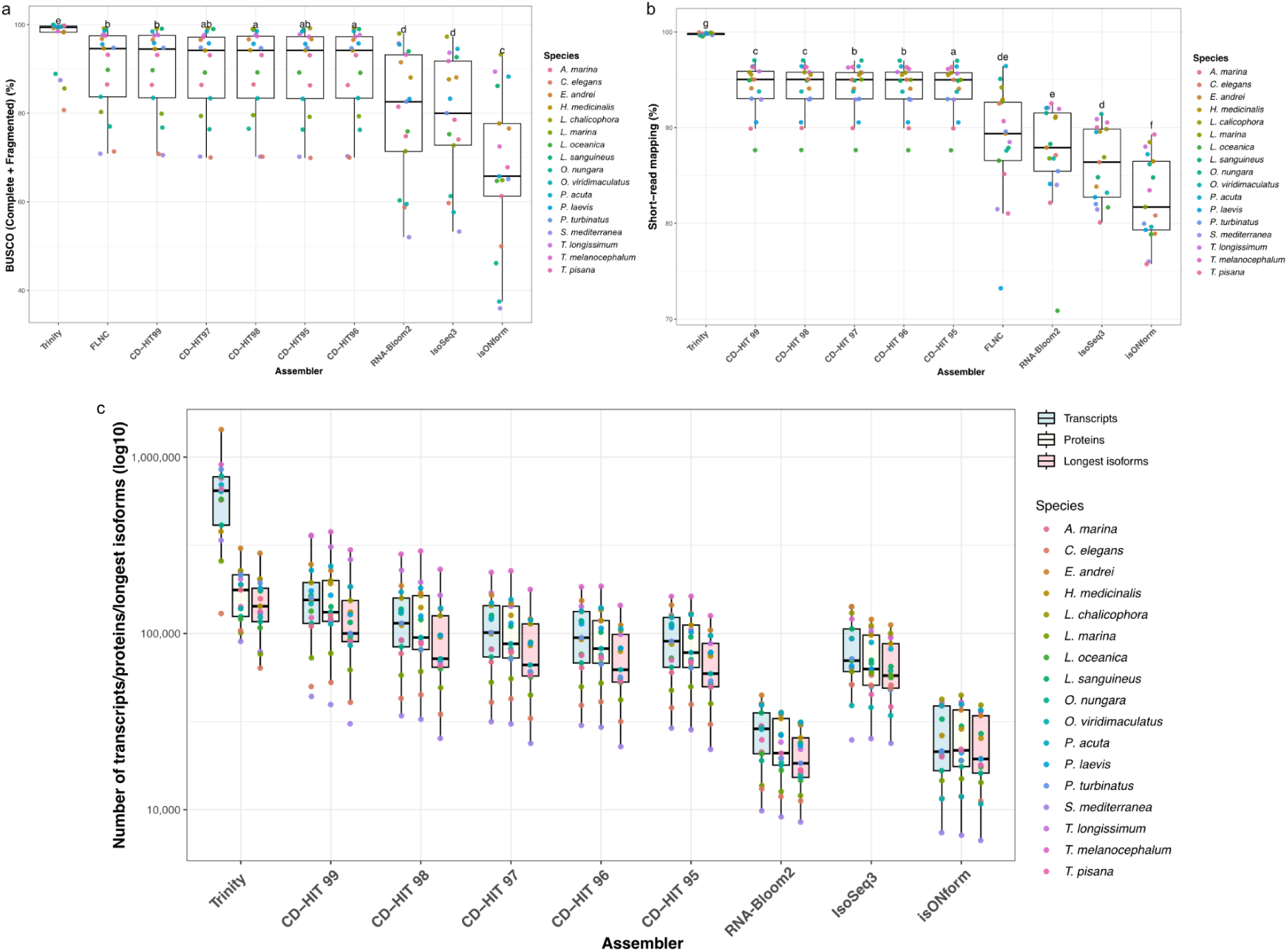
Comparative performance of transcriptome assembly methods across 17 animal species. **(a)** BUSCO completeness scores (metazoa_odb10 database) representing the sum of complete and fragmented single-copy orthologs (C + F%) per assembly. Assemblers are ordered by mean rank across species (Friedman test; post-hoc pairwise Wilcoxon signed-rank tests with Holm correction). Compact letter display (CLD) above each box indicates statistically homogeneous groups (α=0.05). **(b)** Percentage of Illumina short-reads mapping back to each assembly, as a measure of representation of high accurate short-reads in each assembly. Ordering follows the same statistical framework as in (a). **(c)** Log_10_-scaled counts of total transcripts, predicted proteins, and longest protein isoforms identified per assembly. FLNC are excluded, since predicted proteins and longest isoforms values are not applicable. In all panels each colored point represents one species. [*High resolution figure can be found here:* [Figure3.pdf](https://drive.google.com/file/d/1s1PSn68aN3rxR-gzWWX-3cF6Q6_fYXOG/view?usp=share_link)]

**Figure 4.**
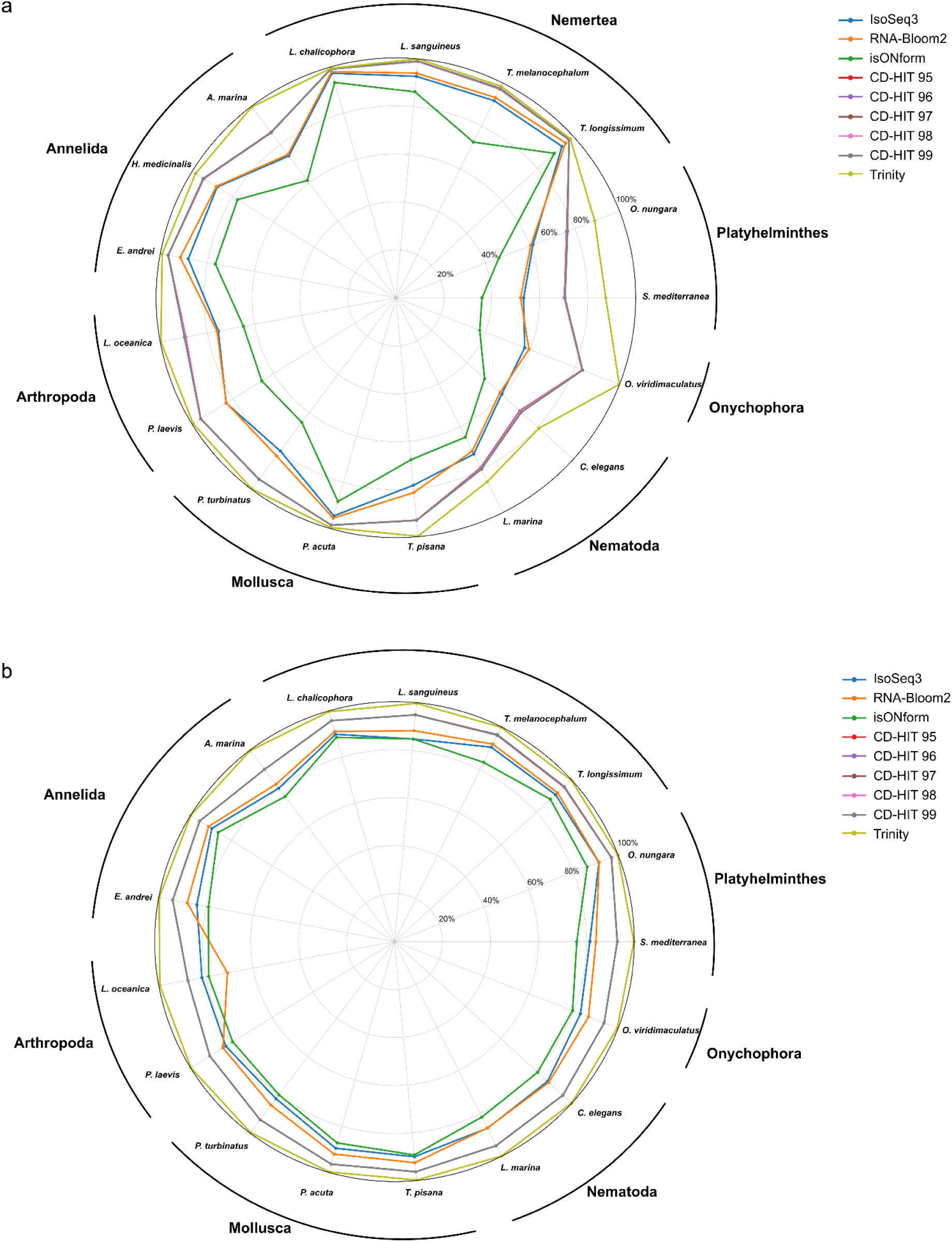
Assemblers performance across species. (a) Radar plot showing BUSCO completeness (Complete + Fragmented) (b) Radar plot showing percentage of short-reads mapped per assembly. [*High resolution figure can be found here:* [Figure4.pdf](https://drive.google.com/file/d/1QgTDRY-frG127E_frkgwDQ2u4isUbFDC/view?usp=share_link)]

Trinity also achieved high completeness but displayed a high rate of duplicated BUSCOs, likely due to transcript redundancy and variability in gene expression levels. These issues are consistent with previously reported limitations of short-read assemblers, including reduced reproducibility and an elevated rate of false positives (Bankar et al. 2015; Wang and Gribskov 2017). In contrast, RNA-Bloom2 produced lower duplication rates (Figure 3a; Figure S1), though at the expense of overall completeness, placing it in a significantly lower CLD group than Trinity and CD-HIT. As expected, unassembled FLNC reads show very high BUSCO duplication, reflecting the presence of multiple overlapping isoforms prior to clustering or assembly. Finally, we note that CD-HIT assemblies also display increased duplication at higher similarity thresholds, illustrating the trade-off between maximizing completeness and minimizing redundancy.

### Read mapping support

Read mapping analyses show that CD-HIT at all similarity thresholds and Trinity assemblies capture the highest proportion of short-reads (Figure 3b, 4b; Figure S2). Trinity’s strong performance is expected, since its assemblies are generated directly from the short-read data that are also used for the mapping. Among the long-read strategies, RNA-Bloom2 consistently achieves higher mapping rates than IsoSeq3 and isONform, indicating that it better represents the transcript diversity detected by Illumina sequencing. In contrast, isONform performs worst across most species, often recovering substantially fewer reads, suggesting that its consensus-based reconstruction discards valid transcript variants present in the data.

### Coding potential

Coding potential and isoform recovery (Figure 3c; Supplementary Figure S3) highlight clear differences among assemblers. Trinity assemblies are highly redundant, producing far more transcripts than unique proteins or isoforms, consistent with earlier reports of transcript fragmentation and over-representation in short-read assemblies (Cerveau and Jackson 2016). In contrast, CD-HIT at 99% similarity generates the largest number of transcripts and efficiently preserves coding information, as most transcripts are retained after translation into proteins.

RNA-Bloom2 produces fewer transcripts and proteins overall, and in many species transcript counts exceed protein counts, suggesting redundancy and the recovery of non-coding sequences. IsoSeq3 and isONform yield more balanced results, with similar counts of transcripts, proteins, and isoforms, reflecting their ability to retrieve full-length isoforms directly from the long-read data.

Published benchmarks of RNA-Bloom2 (Nip et al. 2020) and isONform (Petri and Sahlin 2023), including comparisons against the Nanopore-based assembler RATTLE (de la Rubia et al. 2022), indicate that consensus-based approaches such as the last step of isONform and IsoSeq3 may under-represent transcript diversity and fail to capture biologically relevant isoforms, observations consistent with our results. Together, these results reinforce that direct clustering with CD-HIT, particularly at high similarity thresholds, recovers a richer and more accurate representation of coding potential than either consensus-based or graph-based assembly strategies.

### Structural validation using SQANTI3

To assess the structural accuracy of transcript reconstructions, we analysed all assemblies generated for *Caenorhabditis elegans* using SQANTI3 against the reference genome and annotation (GCF_000002985.6) (Figure 5). SQANTI3 classifies transcripts into four main categories: full-splice match (FSM; incomplete splice match (ISM); novel-in-catalogue (NIC); and novel-not-in-catalogue (NNC). This analysis complements the BUSCO, mapping-rate, and coding-potential assessments by measuring the extent to which reconstructed transcripts correspond to annotated splice structures.

**Figure 5.**
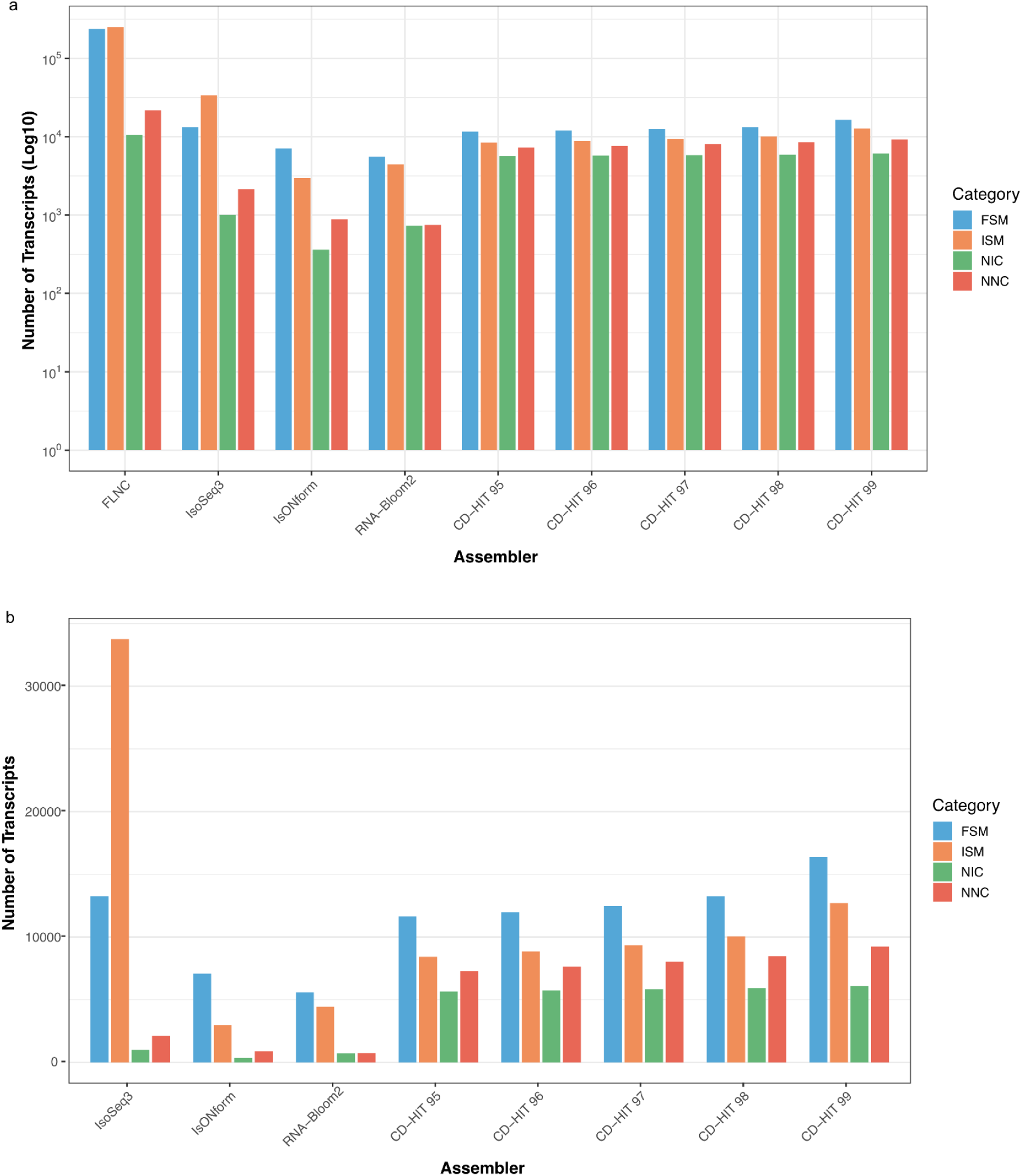
SQANTI3 structural category classification across assemblies. **(a)** Overview of all data including unassembled FLNC reads (log_10_ scale). **(b)** Detailed comparison of assembled transcriptomes only, excluding FLNC (linear scale). Categories: FSM, full splice match; ISM, incomplete splice match; NIC, novel in catalogue; NNC, novel not in catalogue. [High resolution figure can be found here: [Figure5.pdf](https://drive.google.com/file/d/1VlIHWWPXbCK9PtTeagnV0II0N5Dfc43q/view?usp=sharing)]

As expected, the unprocessed FLNC reads contained the largest total number of transcripts across all structural categories, reflecting the redundancy of reads prior to any clustering or assembly step (Figure 5a). Among assemblers, CD-HIT 99 recovered the highest number of FSM transcripts indicating greater structural concordance with known splice junctions, while IsoSeq3 showed comparably high FSMs but disproportionately elevated ISMs, suggesting that it reconstructs partial isoforms rather than full-length transcripts (Figure 5b). CD-HIT assemblies showed a progressive increase in FSMs and ISMs at higher similarity thresholds, probably due to the retention of more complete and redundant isoforms. In contrast, isONform and RNA-Bloom2 yielded substantially fewer transcripts across all categories (Figure 5b). NIC and NNC transcripts were comparatively low across all assemblers, though CD-HIT assemblies recovered more novel transcripts than other methods, with counts increasing at higher similarity thresholds.

Overall, the SQANTI3 analysis corroborates the conclusions obtained from BUSCO completeness and coding-sequence recovery. So far, CD-HIT 99 emerges as the long-read strategy providing the most comprehensive representation of transcript diversity, while retaining known gene models.

## Conclusions

The core aim of this project was to evaluate how effectively transcriptomes can be reconstructed from long-read data de novo, without relying on genomic resources. This is a common scenario in non-model organism research, where high-quality genomes are often unavailable or cost-prohibitive to generate, representing a bottleneck in long-read-based comparative genomics. By comparing de novo assembly and clustering tools across a diverse set of animal lineages, this work defines what can be achieved under these constraints and identifies which approaches yield the most complete and biologically informative transcriptomes in a fully de novo context.

We found that direct clustering with CD-HIT at high similarity thresholds offers the most complete and coding-rich transcriptomes from Iso-Seq data, consistently outperforming consensus-based methods and approaching the completeness of Trinity. Unlike Trinity, CD-HIT avoids excessive redundancy and better preserves the full-length isoforms captured by long reads. Structural classification with SQANTI3 further supports this conclusion by showing that CD-HIT 99 recovered the highest proportion of full splice match transcripts among long-read strategies, confirming not only quantitative completeness but also structural accuracy at the level of known splice junctions.

Our results highlight the practical value of clustering approaches for laboratories working without genomic resources, where reference-guided strategies are not feasible. In this context, CD-HIT provides a realistic and efficient solution for reconstructing transcriptomes from long-read sequencing. At the same time, the comparison to short-read Trinity assemblies underscores the trade-offs between completeness and redundancy that remain central to transcriptome reconstruction regardless of sequencing technology.

## Supplementary Data

Supplementary figures can be found: [Supplementary_Figures.pdf](https://drive.google.com/file/d/1KaGGXmLBmcXR1hMu9Pnp2_5mR-54w5-N/view?usp=sharing)

Supplementary tables can be found here: [Supplementary Tables](https://docs.google.com/spreadsheets/d/1ycfFTF7eYPwouKqG2nJX5PHMRR7PlOiyRdLLSoleXs0/edit?gid=0#gid=0)

## Acknowledgements

R.F acknowledges support from the European Research Council (grant agreement no. 948281). We thank Centro de Supercomputación de Galicia (CESGA) for access to computer resources. ChatGPT was used to polish grammar and improve readability of the manuscript. No AI tool was used for idea generation, data analysis or interpretation of results.

## Code Availability

All scripts used in this study are publicly available in https://github.com/MetazoaPhylogenomicsLab/Eleftheriadi_et_al_2026_Isoseq_evaluation/tree/main

## Author Contributions

K.E. and R.F. conceived and designed the study. K.E. and M.V.-V. performed the data analyses. K.E. and M.V.-V. wrote the first draft of the manuscript. R.F. supervised the research, contributed to the interpretation of the results, and provided critical revisions and feedback on the manuscript. All authors reviewed, approved, and contributed to the final version of the manuscript.

## Conflicts of Interest

The authors declare no conflict of interest.

## Notes

### Competing Interest Statement

The authors have declared no competing interest.

### Summary of Updates

Incorporated new analyses to make our results more robust.

